# Sfp1 regulates the SAGA component Tra1 in response to proteotoxic stress in *Saccharomyces cerevisiae*

**DOI:** 10.1101/384602

**Authors:** Yuwei Jiang, Matthew D. Berg, Julie Genereaux, Khadija Ahmed, Martin L. Duennwald, Christopher J. Brandl, Patrick Lajoie

## Abstract

Proteotoxic stress triggers transcriptional responses that allow cells to compensate for the accumulation of toxic misfolded proteins. Chromatin remodeling regulates gene expression in response to the accumulation of misfolded polyQ proteins associated with Huntington’s disease (HD). Tra1 is an essential component of both the SAGA/SLIK and NuA4 transcription co-activator complexes and is linked to multiple cellular processes associated with misfolded protein stress, including the heat shock response. Cells with compromised Tra1 activity display phenotypes distinct from deletions encoding components of the SAGA and NuA4 complexes, indicating a potentially unique regulatory role of Tra1 in the cellular response to protein misfolding. Here, we employed a yeast model of HD to define how the expression of toxic polyQ expansion proteins affects Tra1 expression and function. Expression of expanded polyQ proteins, mimics deletion of SAGA/NuA4 components and results in growth defects under stress conditions. Moreover, deleting genes encoding SAGA and, to a lesser extent, NuA4 components exacerbates polyQ toxicity. Also, cells carrying a mutant Tra1 allele displayed increased sensitivity to polyQ toxicity. Interestingly, expression of polyQ proteins also upregulated the expression of *TRA1* and other genes encoding SAGA components, revealing a feedback mechanism aimed at maintaining Tra1and SAGA functional integrity. Moreover, deleting the TORC1 (Target of Rapamycin) effector *SFP1* specifically abolished upregulation of *TRA1* upon expression of polyQ proteins. While Sfp1 is known to adjust ribosome biogenesis and cell size in response to stress, we identified a new role for Sfp1 in the control of Tra1, linking TORC1 and cell growth regulation to functions of the SAGA acetyltransferase complex during misfolded protein stress.

## INTRODUCTION

Eukaryotic cells need to correctly fold proteins to ensure their accurate function and avoid the aggregation of toxic misfolded intermediates, which form the basis of several human diseases^1–4^. In Huntington’s disease (HD) expansion of a polyglutamine region encoded by the first exon of the gene encoding the Huntingtin protein (Htt^ex1^) leads to Htt misfolding and aggregation in detergent-insoluble, amyloid-like inclusion bodies (IBs) in the cytoplasm and nuclei of neuronal cells^5–7^. In response to the accumulation of misfolded proteins, including polyQ huntingtin, cells modify their gene expression profile to favor adaptive responses directed at restoring protein homeostasis^8–10^. Well-characterized responses to proteotoxic stress, such as the unfolded protein response of the endoplasmic reticulum^11– 17^ and the heat shock response^18–21^ in the cytoplasm, increase the folding capacity of their respective compartments upon accumulation of misfolded polyQ expansions. These responses prevent the protein quality control machinery from being overwhelmed by sudden changes in the misfolded protein burden.

It is now clear that multiple signaling pathways act in parallel to regulate gene expression during misfolded protein stress. Acetyltransferase complexes regulate chromatin remodeling, a process affected in HD^22–27^. The SAGA (Spt-Ada-Gcn5-Acetyltransferase) and NuA4 (Nucleosome acetyltransferase of H4) complexes were first identified in yeast as containing the lysine acetyltransferases Gcn5 and Esa1, respectively^28–30^. Both complexes have homologues in mammalian cells, hSAGA and Tip60, respectively. The PIKK family member Tra1/TRRAP is an essential component of both SAGA and NuA4 complexes in yeast and mammalian cells^31,32^. The group of PIKK proteins also includes mTOR, ATM and ATR, which are characterized by a C-terminal PI3K domain^33^. Tra1’s role in SAGA and NuA4 is to interact with transcriptional activators thereby recruiting the complexes to target promoters^34–37^. Because of its presence in both SAGA and NuA4, reducing Tra1 function affects cells distinctly from deletions of components specific to either individual complex. For example, impaired Tra1 function causes generation-dependent telomere shortening, a phenotype that is not detected in cells carrying deletions of either SAGA or NuA4 components^38^.

Misfolded polyQ expansions specifically alter the composition of the SAGA complex and SAGA-regulated gene transcription in both yeast and mammalian models^39–44^. These studies employed polyQ-expanded ataxin-7/Sca7, which is responsible for the neurodegenerative disease spinocerebellar ataxia 7^45^. Sca7/ataxin-7 is a component of SAGA and SLIK (SAGA-like) acetyltransferase complexes and thus explaining the effect of polyQ expandedSca7 on SAGA function ^39,40^. Targeting Htt^ex1^ to the yeast nucleus also alters transcription similarly to cells carrying deletions in genes encoding SAGA components^46^; however, the specific molecular mechanism by which Htt^ex1^ polyQ expansions affect SAGA function remains unclear. Our previous genetic screen for synthetic interactions linked Tra1 to the regulation of several stress responses, including protein misfolding stress^47^. Tra1 is therefore a strong candidate target for polyQ proteins to regulate the transcriptional response to protein misfolding stress.

To study the effect of Htt^ex1^ polyQ expansion on yeast, we employed a well-characterized model that involves expressing fluorescently-tagged Htt^ex1^ ^48–52^. We define the interplay between Tra1 and polyQ-induced stress. We also identify the TORC1 effector Sfp1 as a regulator of polyQ toxicity that regulates Tra1 expression during proteotoxic stress, thus expanding our understanding of its role beyond the regulation of cell growth and ribosome biogenesis^53–55^. Our findings further define the roles of TORC1, Sfp1 and Tra1 in response to polyQ proteins.

## MATERIAL AND METHODS

### Yeast genetic manipulation and growth assays

All strains are derivatives of either BY4741/4742 or W303a (**see Supplemental Table 1**). Gene deletions were performed using standard yeast genetics procedures^56^ and validated by sequencing. Plasmids were transformed using the lithium acetate method^57^. Cell growth was assessed by both spot assay on agar plates and growth in liquid culture. Yeast cells were cultured overnight in selective synthetic complete media with 2% glucose as a sole carbon source. For spot assays, cultures were diluted to equal concentrations and then spotted in 4 fivefold dilutions using a pinning tool with the most concentrated spot equalized at OD_600_0.2. Cells were grown on selective plates at 30°C for 2 days and imaged using a Geldoc system (Bio-RAD). For liquid culture, cells were diluted to OD_600_ 0.1 and incubated at 30°C. OD_600_ was measured every 15 min using a BioscreenC plate reader (Growth curves USA) for 24 hours. Growth curves were generated and the area under the curve calculated for each biological replicates and a two-tailed student t-test was used to determine statistical significance between the different experimental conditions using Graphpad (Prism).

### Drugs

Stock solutions of tunicamycin (5 µg/ml in DMSO; Amresco), Trichostatin A (10 mM in H_2_O; Biovision), calcofluor white (30 mg/ml in H_2_O; Sigma-Aldrich), rapamycin (1 mg/ml in DMSO), H_2_O_2_ (9.79 M) cycloheximide (10 mg/ml in water) (Fisher Scientific), and MMS (99%; Acros Organics) were prepared and used at the indicated concentrations.

### DNA constructs

Plasmids encoding fluorescently tagged Htt^ex1^ and LacZ reporter constructs carrying *TRA1, PHO5, and PGK1*^58,59^ promoters in YCplac87^60^ were previously described (**see Supplemental Table 2**). *SPT7* promoter sequences relative to the translational start, -633 to +68, *NGG1* promoter sequences -430 to +5 and *EAF1* promoter sequences -890 to +31 were engineered by PCR as BamHI/HindIII fragments using oligonucleotides listed in Table 3 and cloned into YCplac87^60^ to generate transcriptional reporters. Vectors encoding fluorescently tagged Tra1 with either ysmfGFP^52^ and yemRFP^61^ were generated by replacing the eGFP coding sequence by the new codon-optimized fluorescent proteins using the BamHI/NotI sites in the previously described eGFP-Tra1 vector^62^ using primers listed in **Supplemental Table 3**.

### Fluorescent microscopy

Cells were diluted 10X and transferred to LabTek imaging chambers (Thermo Inc.) and imaged at room temperature. Fluorescent microscopy was performed using a Zeiss 800 confocal microscope equipped with a 63× PlanAprochromoat objective (1.4 NA). Images were analyzed using the ImageJ software^63^.

### qRT-PCR

RNA extraction was performed using MasterPure Yeast RNA Purification Kit (Lucigen). cDNA synthesis was done by qScript Flex cDNA Synthesis Kit (Quanta Bioscience). The cDNA preparations were used as the template for amplification using PerfeCTa SYBR-Green Supermix (Quanta Bioscience). The primers used were listed in supplemental Table 3. The relative expression level was calculated using the comparative Ct method and U3 was used as a reference gene.

### Western blot

Yeast cells were lysed using 0.1 M NaOH for 5 mins at room temperature, resuspended in SDS sample buffer and boiled for 5 mins^64^. Proteins were separated using gel electrophoresis and transferred to PVDF membrane. The membrane was blocked with 5% milk. Then the membrane was incubated with anti-Flag (M2, Sigma-Aldrich), anti-PGK1 (Invitrogen), anti-histone H3 and anti-histone H3K14 (Abcam) overnight, followed by 1 h incubation with the appropriate fluorescent secondary antibody and imaged with an Odyssey infrared imager (Licor) to detect the signal.

### β-galactosidase assay

Cells were harvested and resuspended in lacZ buffer. To measure β -galactosidase activity, 50 µl cell lysate was mixed with 950 µl lacZ buffer containing 2.7 µl β-mercaptoethanol, 1 drop 0.1%SDS, 2 drop CHCl_3_ and incubated at 30°C for 15 min. The reaction was started by adding 100 µl ONPG (4 mg/ml) and incubated at 30°C till the color changed to yellow. 300 µl 1 M Na_2_CO_3_ was added to stop the reaction. The β-galactosidase activity was determined at 420nm absorbance using a plate reader, normalizing data to cell density.

### Data availability

All strains and plasmids are available upon request. The authors affirm that all data necessary for confirming the conclusions of the article are present within the article, figures and tables.

## RESULTS

### PolyQ expansions compromise the SAGA acetyltransferase complex

In our experiments, Htt^ex1^ is placed under the control of the *GAL1* promoter and induced by growth n galactose as sole carbon source. Under these conditions, expressing HD-associated polyQ lengths (46Q and 72Q) results in a polyQ length-dependent growth defect compared to the non-HD associated 25Q^14,50,51^ (Figure 1A). As opposed to what is observed for other disease-causing misfolded proteins, such as α-synuclein, polyQ expression inhibits cell growth but does not cause significant cell death as measured by either regrowth assays or labeling of dead cells with propidium iodide (Supplemental Figure 1). The effect of a 103Q Htt^ex1^ polyQ expansion is also apparent when expressed at more modest levels under the transcriptional control of the relatively weak *MET25* promoter^14^ (Figure 1B). This model allows testing low polyQ toxicity without altering the carbon source. High expression of polyQ expanded Htt^ex1^ results in polyQ length-dependent formation of cytoplasmic aggregates that can be observed using fluorescent microscopy (Figure 1C).

**Figure 1:**
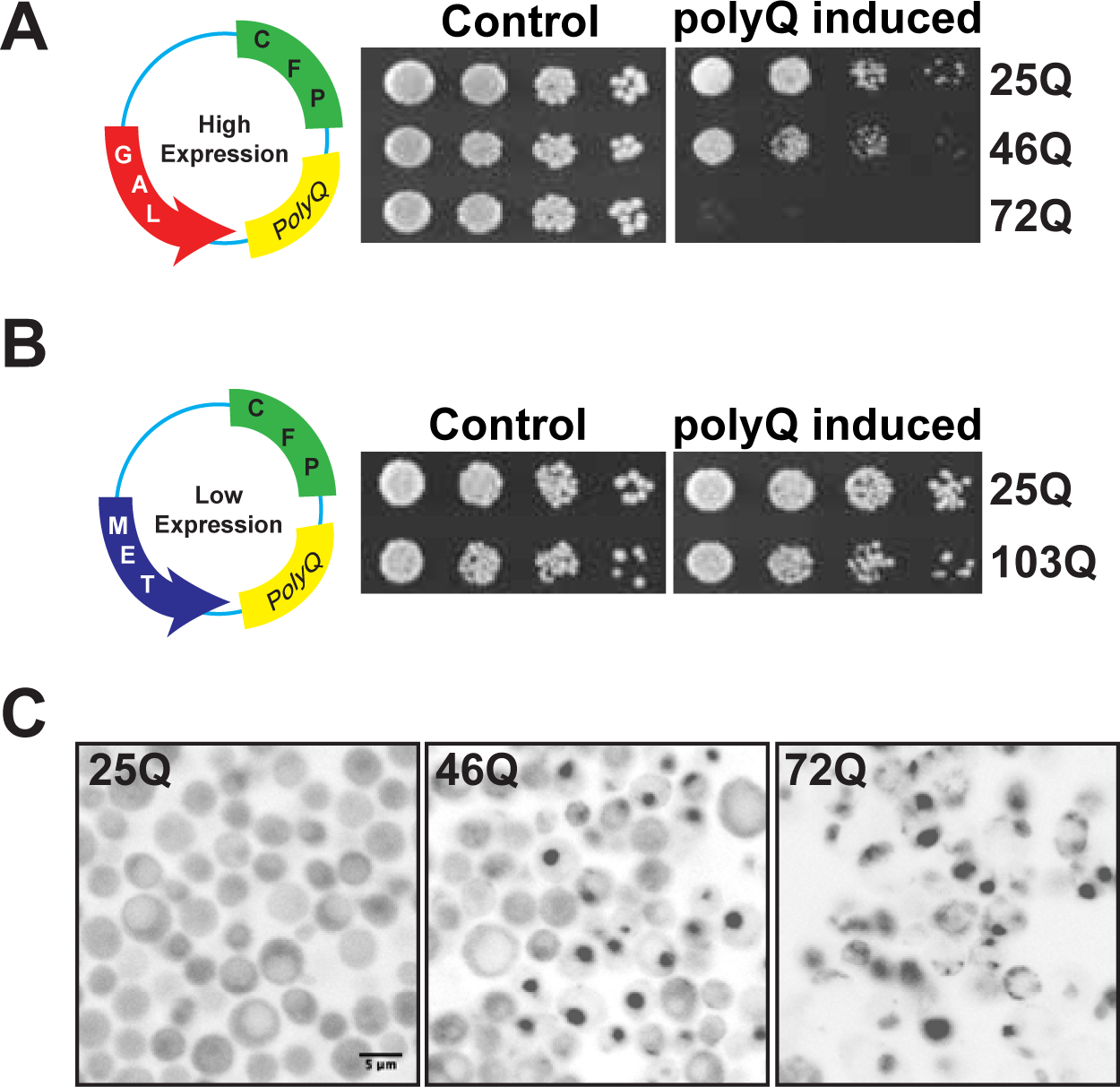
PolyQ toxicity and aggregation in yeast. **(A)** The yeast model of HD. Cell expressing high levels of CFP-tagged Htt^ex1^ display polyQ length-dependent toxicity. 25Q serves as a control for non-pathological Htt^ex1^. Cell growth was assessed by serial dilutions on SC plates containing either glucose (control) or galactose (polyQ induced). (**B**) Cells expressing low levels of CFP-tagged Htt^ex1^ display only modest growth defect even in presence of 103Q. Cell growth was assessed by serial dilutions on SC plates containing presence (control) or absence (polyQ induced) of methionine. (**C**) Fluorescent images show accumulation of inclusion bodies in 46Q and 72Q-expressing cells as opposed to diffused cytosolic distribution of 25Q after induction in galactose.

Misfolded polyQ proteins associated with the polyglutamine disease spinocerebellar ataxia disrupt assembly of the SAGA complex and SAGA-dependent transcription^40^. The ensuing phenotype resembles those associated with deletions of SAGA complex components. Sca7, the protein responsible, is a subunit of the hSAGA complex. Thus, its crucial role in SAGA function is expected. To investigate the relationship between Htt^ex1^, a protein that is not part of the acetyltransferases complexes, we examined genetic interactions of Htt^ex1^ with deletions of NuA4 (*eaf1Δ, eaf6Δ, yaf9Δ*) and SAGA (*ada2Δ, spt8Δ, ubp8Δ*) components (Figure 2A). Expanded polyQ expression displayed increased growth defects with deletion of the SAGA components Ada2 and Ubp8, but not Spt8. 72Q expression only modestly affected growth in one of the NuA4 related deletions, *eaf6Δ*. For Tra1, we employed a previously characterized loss-of–function mutant of *TRA1* (*tra1-F3744A*) that carries a mutation in the C-terminal FATC domain^65^. *tra1-F3744A* displayed strong growth defects in presence of expanded 72Q in both plate and liquid growth assays (Figure 2B). Supporting these data, polyQ expression sensitized cells to stresses that result in slow growth of strains carrying *TRA1* mutations, i.e. calcofluor white, growth at 39°C, 5% ethanol, MMS and caffeine^59^ (Figure 2C). Expression of expanded polyQ proteins was also associated with decreased activity from SAGA regulated promoters (*HIS4* and *PHO5*) (Figure 2D) and histone H3 acetylation (Figure 2E), implicating that polyQ affects SAGA function in yeast.

**Figure 2:**
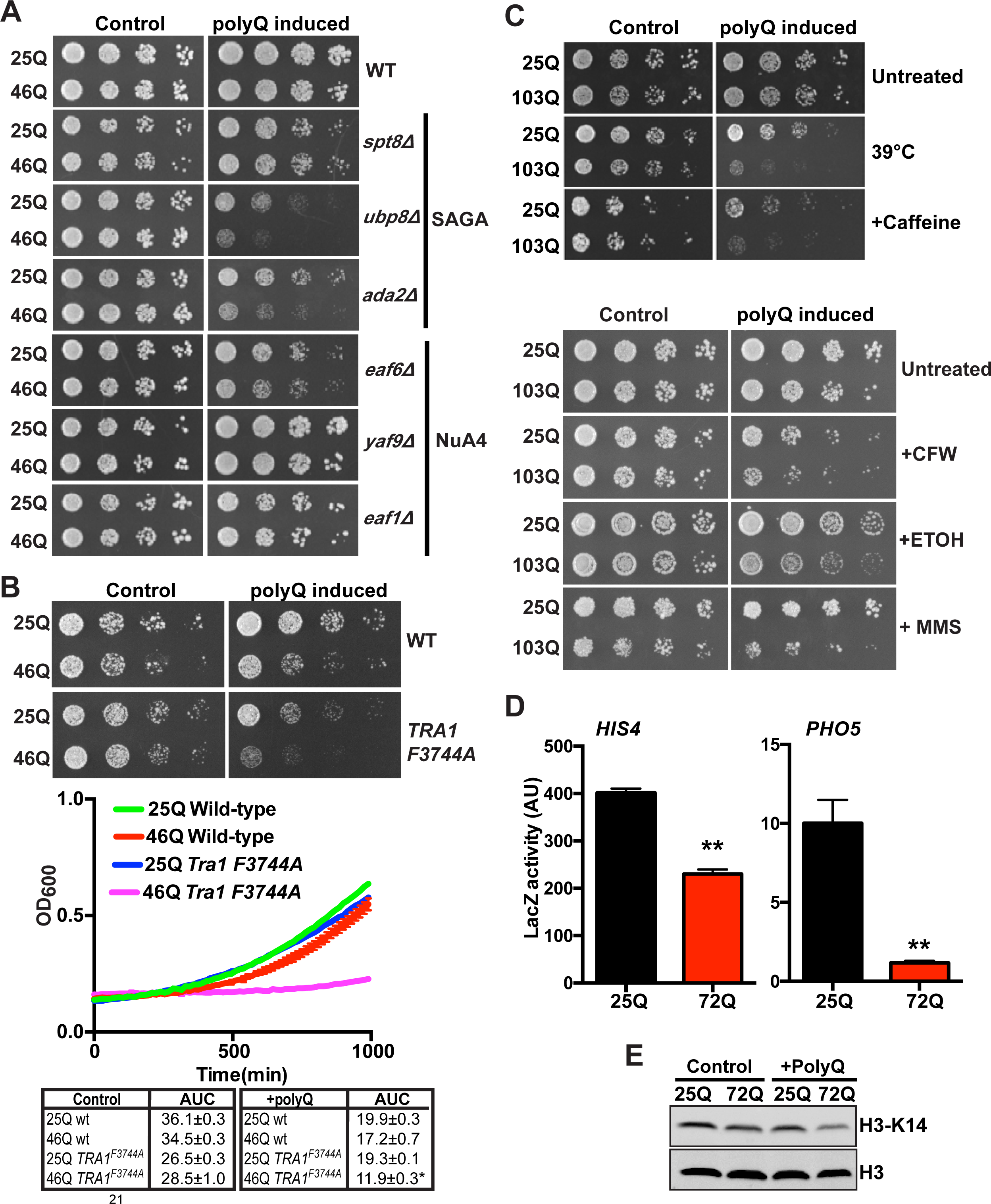
polyQ expression and SAGA and NuA4. **(A)** Wild-type, *ada2Δ, spt8Δ, ubp8Δ* (SAGA) and *eaf1Δ, eadf6Δ, yaf9Δ* (NuA4) cells expressing either 25Q or 46Q Htt^ex1^ assessed by growth assays on SC media plates in presence of glucose (control) or galactose (polyQ induced). **(B)** Wild-type and *tra1-F3744A* cells expressing either 25Q or 46Q Htt^ex1^ were assessed by growth assay on SC media plates and in liquid growth assay in presence of glucose (control) or galactose (polyQ induced). The area under the curve was quantified for each replicates (n=3) **p<0.005. **(C)** Cells expressing either 25Q or 103Q Htt^ex1^ were spotted on plates containing various stressors (0.1% caffeine, 6µg/ml calcofluor white (CFW), 5% ethanol (ETOH), 0.1% caffeine, 0.03% methyl methanesulfonate (MMS) or incubated at 39°C) in presence or absence of methionine. **(D)** Expression of 72Q induced a significant decreased in the expression from the SAGA-regulated genes *HIS4 and PHO5*. Gene expression was analyzed using LacZ transcriptional reporters was assessed after overnight induction of polyQ in galactose under conditions that induced each promoter (absence of histidine or low phosphate respectively). **p<0.005, n=3±SEM. **(E)** Reduced histone acetylation in cells expression expanded polyQ. Immunoblot of total (H3) and acetylated histone H3 (H3-K14) with or without expression of 25 and 72Q.

### Accumulation of misfolded polyQ expansions increase *TRA1* expression

We previously observed that Tra1 bearing mutations in its PI3K domain (*tra1_Q3_*) decrease nuclear localization of the protein^59^. We therefore assessed whether expressing polyQ expansion affects Tra1 localization using confocal microscopy. We found that expanded polyQ expression (72Q-ymsGFP) did not change the nuclear localization of fluorescently tagged Tra1 (yemRFP-Tra1) (Figure 3). Consequently, polyQ possibly alters SAGA function independently of Tra1 sequestration into cytoplasmic polyQ inclusions. Cells also respond to the defective *tra1_Q3_* allele by increasing the transcription of the *TRA1* gene^59^. We assayed expression from the *TRA1* promoter to evaluate whether expanded polyQ results in a Tra1 defect that also feedbacks to increase *TRA1* expression. As shown in Figure 4A, we found a ~2.5 fold increase in *TRA1* expression of a *TRA1-*promoter LacZ fusion upon expression of 72Q compared to the non-toxic 25Q. Increased transcription was also observed from promoters of other components of SAGA and NuA4, i.e. *NGG1*, *SPT7* and *EAF1*, but not for the control gene *PGK1* as was the case for *tra1*^59^ (Figure 4A). We also observed that the polyQ-induced increase in *TRA1* expression was abolished when cells expressing 72Q were treated with the HDAC inhibitor trichostatin A (TSA) (Figure 4B), suggesting that changes in chromatin remodeling the regulate *TRA1* expression in the presence of polyQ expansion. As a consequence of increased mRNA levels, Tra1 protein abundance increased upon expression of 72Q (Figure 4C). These results suggest that expanded polyQ impairs Tra1 function inducing a feedback mechanism to cope with disrupted SAGA/NuA4 function. As shown in Figure 4D, the increased *TRA1* expression was specific to polyQ and was not observed with other stressors that result in protein misfolding, such as induction of endoplasmic reticulum stress by tunicamycin, heat shock, oxidative stress caused by H_2_O_2,_ or perturbation of cell wall integrity by calcofluor white (Figure 4D). Moreover, expression of α-synuclein, another disease causing misfolded protein, reduced *TRA1* promoter activity, indicating that various misfolded proteins differentially impact *TRA1* transcriptional regulation (Figure 4E). Interestingly, disruption of the SAGA/NuA4 complex activity by deleting *GCN5*, *SPT20* or *EAF3* did not cause *TRA1* upregulation (Figure 4F). These results suggest that the upregulation of *TRA1* by misfolded polyQ is not solely a consequence of disruption of the SAGA complex.

**Figure 3:**
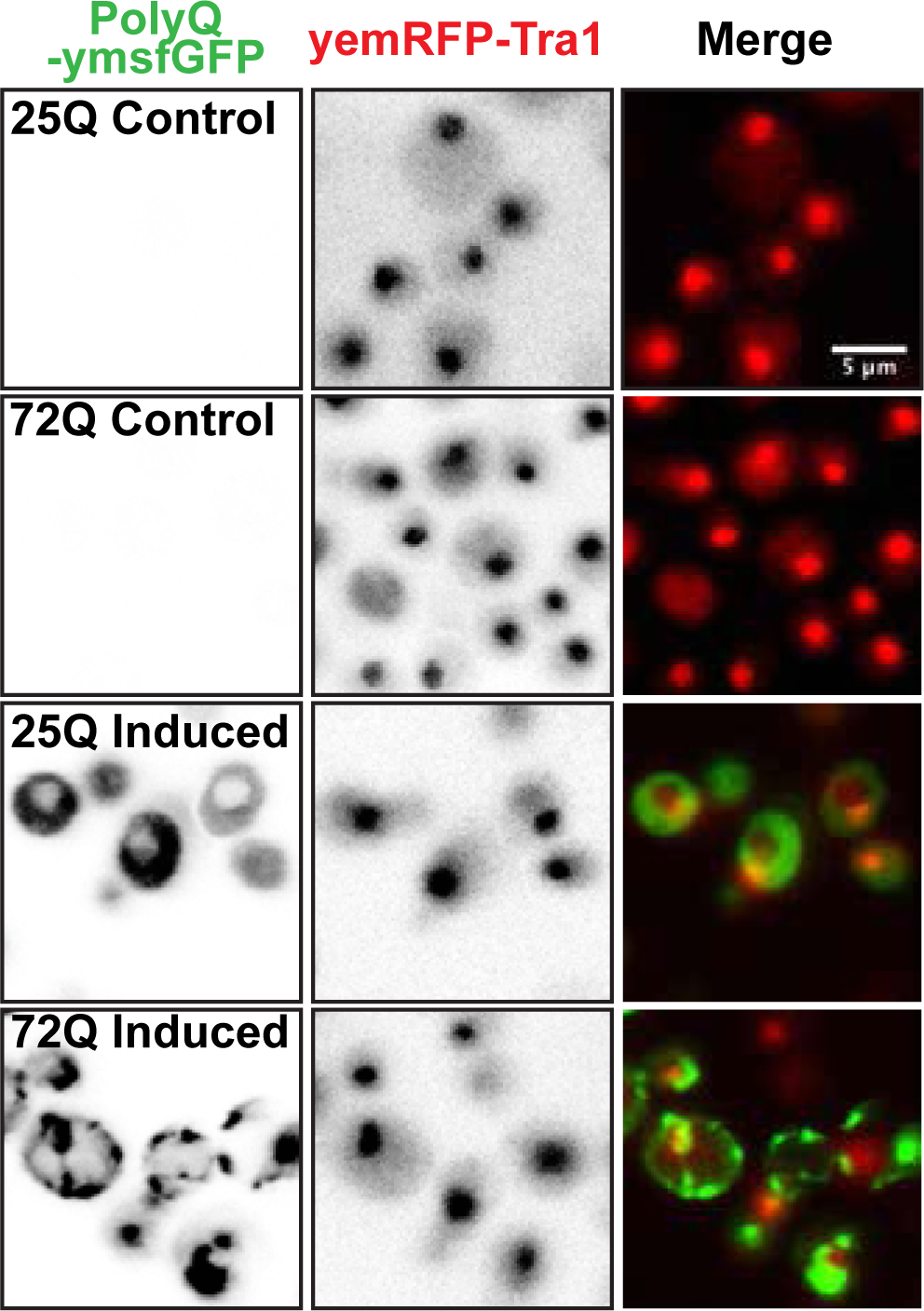
Tra1 is not sequestered into polyQ inclusion bodies. Tra1 remains localized to the nucleus in the presence of polyQ aggregates. Fluorescent images showing localization of polyQ-ymsfGFP and yemRFP-Tra1 after overnight culture in glucose (control) or galactose (polyQ induced).

**Figure 4:**
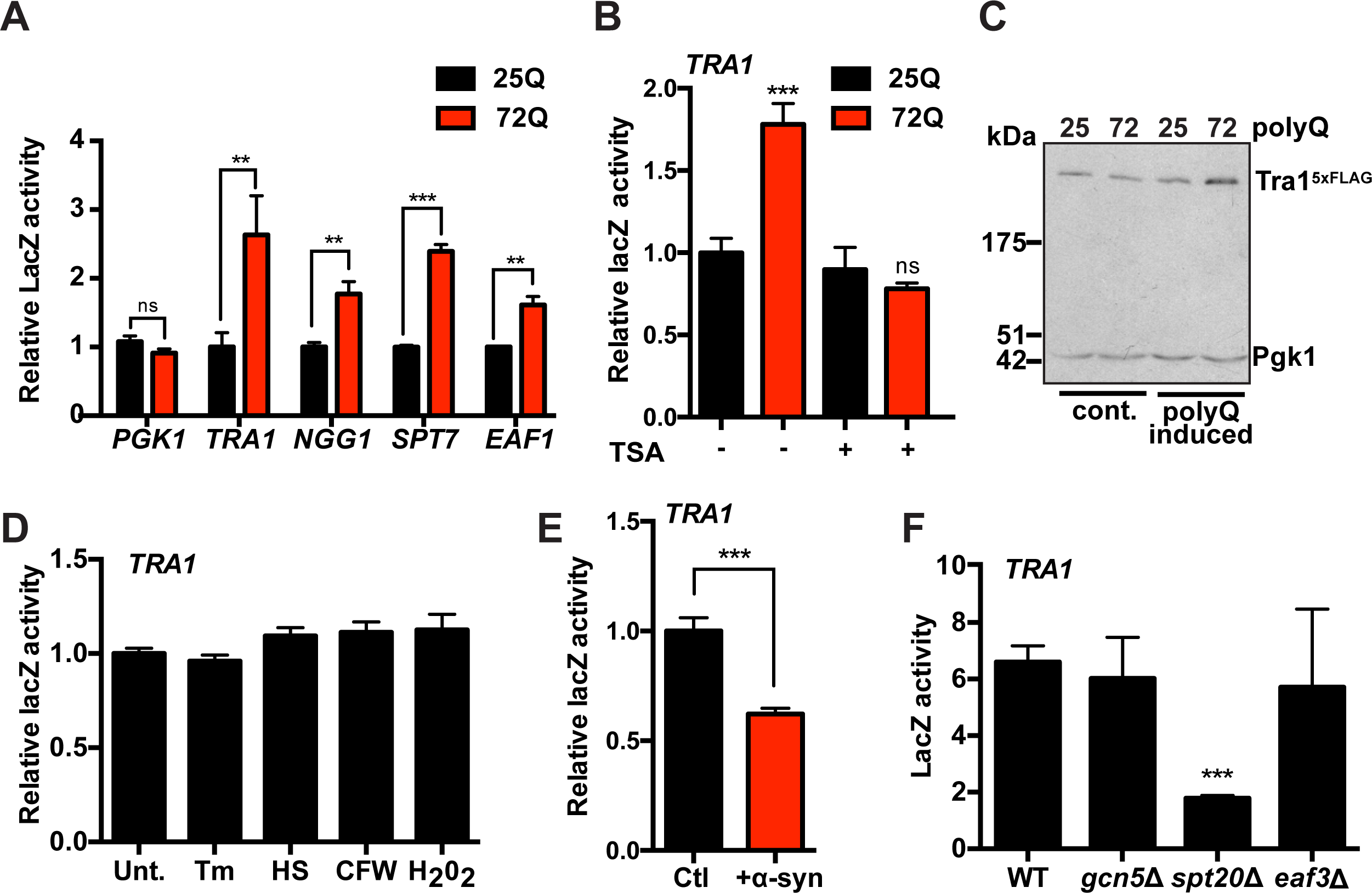
*TRA1* expression is increased in presence of toxic polyQ expansions. **(A)** Increased expression of *TRA1* and other SAGA (*NGG1*, *SPT7*) and NuA4 (*EAF1*) components after overnight induction of 72Q-CFP compared to 25Q-CFP. Expression was analyzed using promoter-LacZ reporters. *PGK1* serves as a control gene. **p<0.005, ***p<0.001 n=3±SEM **(B)** Treatment with the HDAC inhibitor trichostatin A (TSA) abolished *TRA1* upregulation caused by polyQ expansions. 25 and 72Q-CFP were induced overnight in galactose media in presence or absence of 80 µM TSA. (**C**) Expanded polyQ increases Tra1 protein abundance. Cells expressing either 25 or 72Q-CFP and a chromosomally integrated Tra1^5x-Flag^ were cultured in glucose or induced overnight in galactose and processed for immunoblot. (**D**) Effects of other stressors on *TRA1* expression. Cells expressing the *TRA1*-LacZ reporter were treated with tunicamycin (Tm; 5 µg/ml), heat shocked at 42°C, treated with calcofluor white (CFW; 300 µg/ml) or H_2_O_2_ (300 µM) for 2 hr. n=3 ±SEM. (**E**) Expression of α-synuclein decreased *TRA1* expression. Cells expressing a galactose inducible version of α-synuclein-GFP were cultured in either glucose or galactose overnight and TRA1 expression was measured using a LacZ reporter. n=3. **(F)** Deletion of genes encoding SAGA components does not lead to *TRA1* upregulation. Wild-type and *gcn5Δ, spt20Δ* and *eaf3Δ* cells expressing the *TRA1*-LacZ reporter were analyzed. β-galactosidase activity is shown as the average of six replicates with the SD indicated by the error bars.

### TOR1 signaling regulates polyQ toxicity

Our previous genetic screen highlighted a potential role for Tra1 in stress responses^47^, including the control of cellular homeostasis by the TORC1 (Target of Rapamycin Complex 1) regulated signaling cascades that link nutrient availability to cell growth and division. In this screen, a *tra1* mutant displayed a synthetic slow growth phenotype with a deletion of *tor1*^47^. Since data in mammalian cells support both a protective and adverse role for TORC1 in HD^66–71^, we tested the effect of modulating TORC1 activity on polyQ toxicity in yeast. First, we determined the effects of TORC1 inhibition using rapamycin (Figure 5A and B). We found that rapamycin treatment significantly reduced growth of cells expressing expanded 103Q protein in both solid and liquid media assays. Interestingly, rapamycin treatment did not increase polyQ aggregation (Figure 5C). This result argues against a protective role for TOCR1 inhibition by rapamycin through stimulating polyQ aggregate removal through autophagy^68^. Similar to rapamycin, a hyperactive allele of *TOR1* (*TOR1^L2134M^*)^72^ also exacerbated polyQ toxicity (Figure 5D and E). The *TOR1* mutant had no effect on the toxicity of TDP-43 (Figure S2), a protein linked to amyotrophic lateral sclerosis (ALS)^73^, indicating that the role of TORC1 may diverge in different diseases. TORC1 hyperactivation did not prevent the formation of polyQ aggregates (Figure 5F). These data indicate that precise regulation of TORC1 signaling is crucial for cells to cope with polyQ expansion. TORC1 regulates translation and ribosome biogenesis in both yeast and mammals. Incidentally, cells expressing 72Q displayed increased sensitivity to the translational inhibitor cycloheximide (Figure 5G) and decreased expression of ribosomal protein genes (Figure 5H) when compared to 25Q, further supporting a role for TORC1 signaling in polyQ toxicity. Therefore, we next investigated the role of downstream TORC1 effectors in polyQ toxicity.

**Figure 5:**
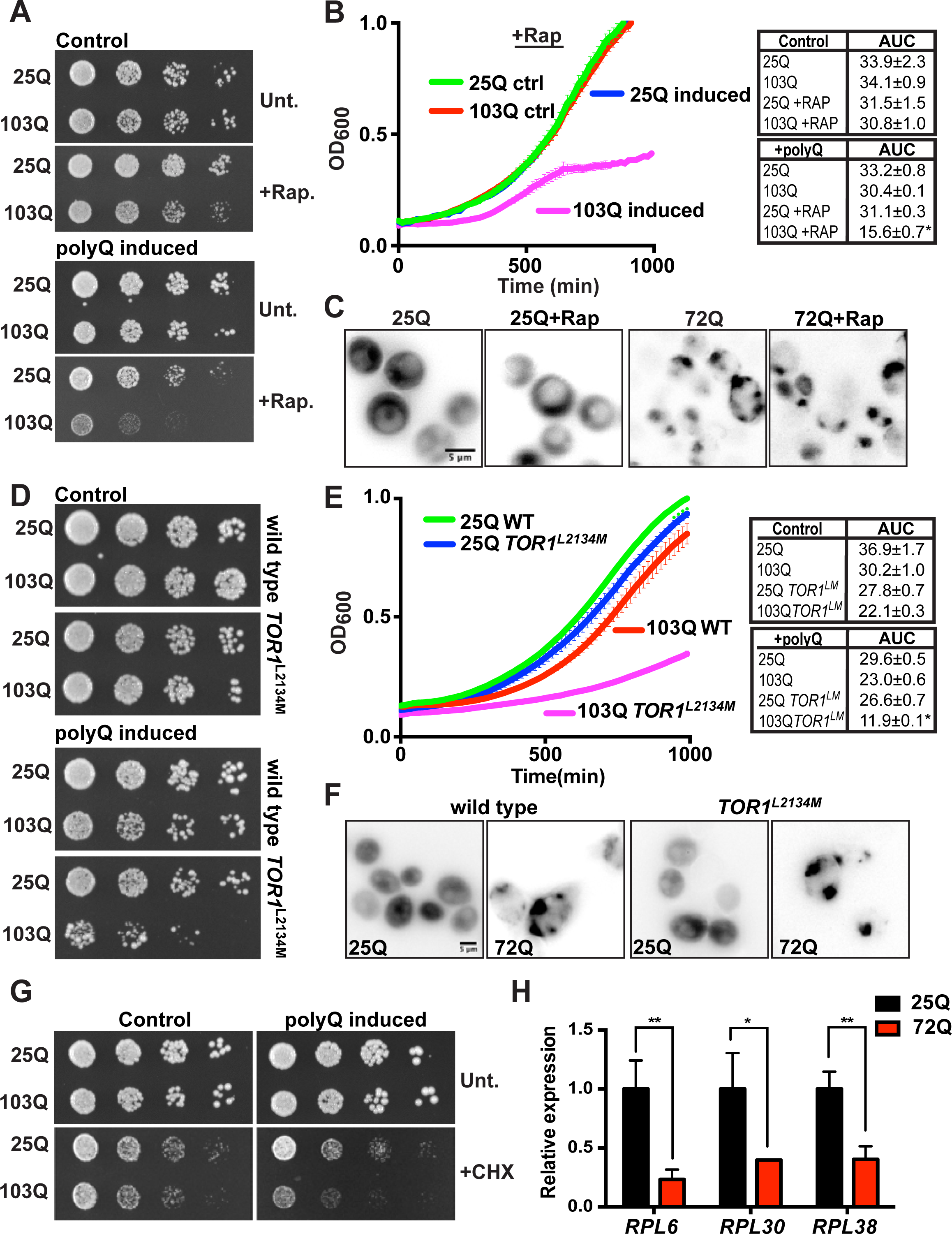
TORC1 regulates polyQ toxicity in yeast. **(A)** TORC1 inhibition by rapamycin exacerbates polyQ toxicity. Wild-type cells expressing either 25Q or 103Q Htt^ex1^ assessed by growth assay on SC containing media plates untreated or supplemented with 2 ng/ml rapamycin in presence (control) or absence (polyQ induced) of methionine. **(B)** Wild-type cells expressing either 25Q or 103Q Htt^ex1^ were cassessed by liquid growth assay in presence (control) or absence (polyQ induced) of methionine ± 2 ng/ml rapamycin. The area under the curve was quantified for each replicates (n=3) ***p<0.001. **(C)** Representative fluorescent images of wild-type cells expressing 25Q or 72Q Htt^ex1^-ymsfGFP after overnight induction in galactose containing media untreated or supplemented with 100 ng/ml rapamycin. **(D)** Wild-type and *TOR1^L2134M^* cells expressing either 25Q or 103Q Htt^ex1^ assessed by growth assay on SC containing media plates in presence (control) or absence (polyQ induced) of methionine. **(E)** Wild-type and *TOR1^L2134M^* cells expressing either 25Q or 103Q Htt^ex1^ were assessed by liquid growth assay in presence (control) or absence (polyQ induced) of methionine ± 100 ng/ml rapamycin. The area under the curve was quantified for each replicates (n=3) ***p<0.001. **(F)** Representative fluorescent images of wild-type and *TOR1^L2134M^* cells expressing 25Q or 72Q Htt^ex1^-ymsfGFP after overnight induction in galactose containing media. (**G**) PolyQ increased cell sensitivity to cycloheximide. Wild-type cells expressing either 25Q or 103Q Htt^ex1^ assessed by growth assay on SC containing media plates untreated or supplemented with 0.1 µg/ml cycloheximide in presence (control) or absence (polyQ induced) of methionine. **(H)** PolyQ expression decreased expression of ribosomal protein genes. RNA was isolated from wild-type cells expressing 25Q or 72Q Htt^ex1^-CFP after overnight induction in galactose containing media and processed for RT-qPCR to assess transcript levels of *RPL6*, *RPL30* and *RPL38*.

### Sfp1/TORC1 regulates *TRA1* expression

In yeast, TORC1 controls gene expression via two main downstream effectors, the mammalian S6 kinase homologue Sch9 and the transcription factor Sfp1^74^. Sch9 localizes to the vacuolar membrane and mediates TORC1 signaling that regulates ribosomal protein gene expression and cell cycle progression^75^. Tor1 also interacts with the transcription factor Sfp1, which regulates expression of ribosomal proteins^54,55,76,77^. During exponential growth, Sfp1 localizes to the nucleus, where it drives transcription of ribosomal protein genes. During various types of stress, including protein misfolding, oxidative stress, and nutrient deprivation, Sfp1 translocates to the cytoplasm^54,55,76,77^ where it is degraded by the proteasome^78^. Here, we found that deleting *SFP1*, but not *SCH9* exacerbated polyQ toxicity (Figure 6A). This agrees with Sfp1 and Sch9 having non-overlapping functions^55^. Indeed the two deletion strains display specific phenotypes. Relevant to our study, *sfp1Δ* increases rapamycin sensitivity as compared to *sch9Δ*^77^. Increased polyQ toxicity in *sfp1Δ* cells was not associated with significant changes in the ability of cells to form inclusion bodies (Figure 6B). Expression of ribosomal protein genes decreased in the presence of misfolded polyQ (Figure 5H) consistent with a loss of Sfp1-mediated TORC1 signaling. In accordance with this, Sfp1-GFP relocalized to the cytoplasm upon 72Q expression, in contrast to its nuclear localization in cells expressing non-toxic 25Q (Figure 6 C and D). It therefore appears that Sfp1 regulates the expression of factors required to cope with expanded polyQ proteins. This may include the global effects on translation due to targeting the ribosomal protein genes, since loss of protein translation due to cycloheximide treatment exacerbates polyQ toxicity. In the same way, reduced translation is linked to polyQ toxicity in mammalian cells^16^.

**Figure 6:**
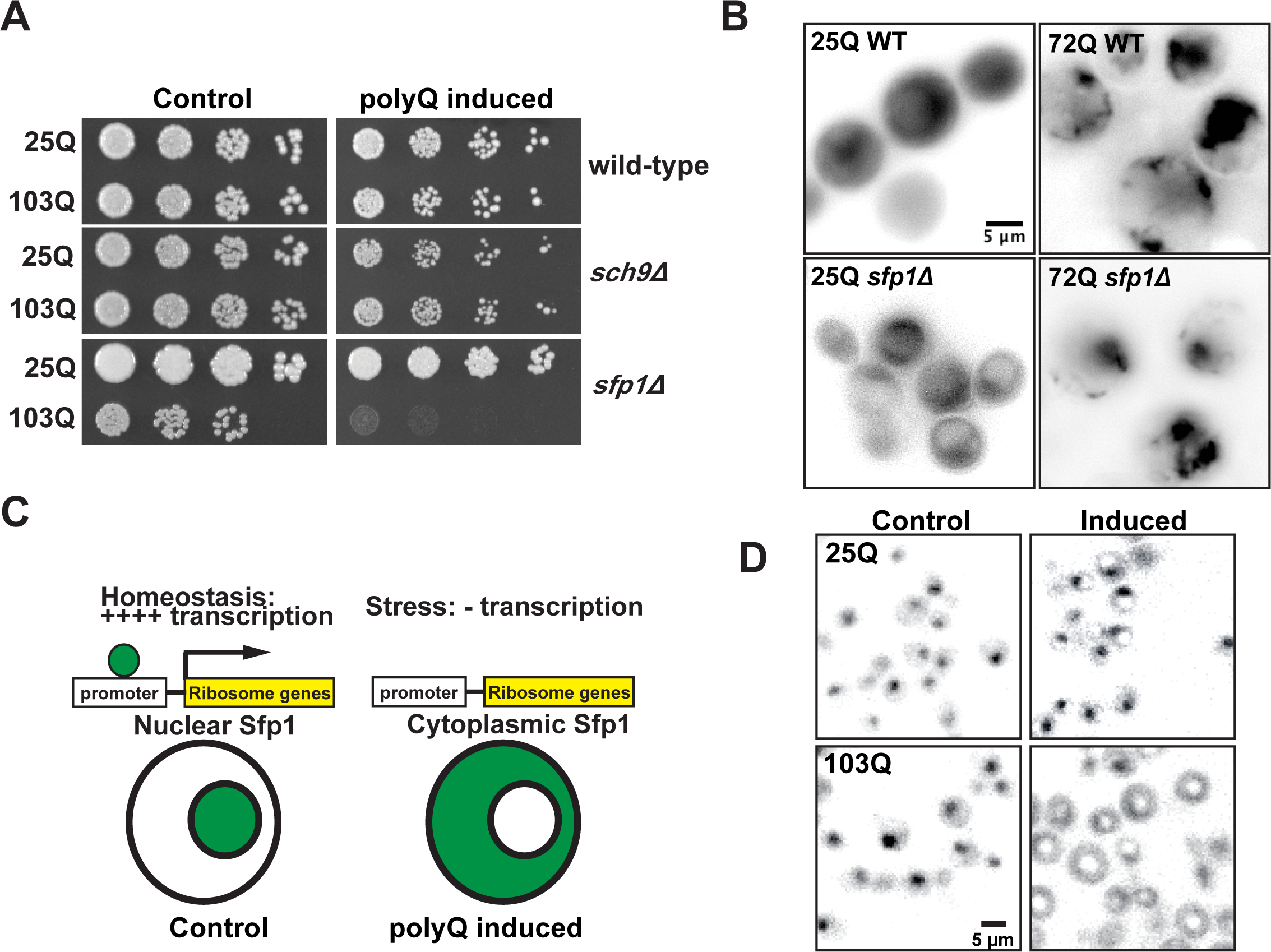
Sfp1 regulates polyQ toxicity. **(A)** Deletion of *SFP1* but not *SCH9* exacerbates polyQ toxicity. Wild-type, *sfp1Δ* and *sch9Δ* cells expressing either 25Q or 103Q Htt^ex1^ assessed by growth assay on SC media plates in presence (control) or absence (polyQ induced) of methionine. **(B)** Deletion of *SFP1* does not affect formation of polyQ IBs. Representative fluorescent images of wild-type and *sfp1Δ* cells expressing 25Q or 72Q Htt^ex1^-ymsfGFP after overnight induction in galactose containing media. **(C)** Diagram that illustrates the relocalization of Sfp1-GFP from the nucleus to the cytoplasm upon stress. The phenotype is linked to decreased sfp1-dependent transcription of RP and RiBi genes. **(D)** Decreased Sfp1 nuclear localization upon expression of expanded polyQ. Representative fluorescent images of wild-type and *sfp1Δ* cells expressing 25Q or 103Q Htt^ex1^-RFP and Sfp1-GFP after overnight induction in galactose containing media.

Interestingly, Lempiäinen *et al*^77^ demonstrated that Sfp1 physically interacts with Tra1, suggesting that Tra1 may be targeted by Sfp1in the presence of PolyQ proteins. As shown in Figure 7A, Tra1 protein levels but not mRNA (Figure 7A and B) were modestly decreased in *sfp1Δ* cells, potentially reflecting reduced global translation due to decreased ribosome production^53–55,77^. Thus, it appears that Tra1 function rather than levels are affected by Sfp1. Indeed, *sfp1Δ* cells are sensitive to high temperature, ethanol, MMS and calcofluor white treatment, hallmarks of impaired Tra1 functions (Figure 7C). Reduced Tra1 activity in *sfp1Δ* cells was further supported by the reduced expression of *PHO5*, a Tra1/SAGA regulated gene^80,81^. This phenotype was reversed by the HDAC inhibitor TSA (Figure 8A), indicating that deleting *SFP1* may affect SAGA and possibly NuA4-mediated chromatin modifications that regulate *PHO5* transcription^80,81^. We next tested the effect of deleting *SFP1* on the polyQ-induced upregulation of SAGA and NUA4 components observed in Figure 2D. Whereas the protein levels of *SPT7*, *NGG1* and *EAF1* were still increased upon expression of misfolded 72Q, *SFP1* deletion abolished *TRA1* upregulation (Figure 8B). These results suggest that Sfp1 specifically regulates *TRA1* expression upon polyQ expression. Sfp1 is a known TORC1 effector and TORC1 inhibition with rapamycin downregulates *TRA1* expression^54^ (Figure 8C) highlighting a role for TORC1 in the control of *TRA1* transcriptional regulation. Since deleting *SFP1* causes hyperactivation of TORC1 as a compensatory mechanism^77,82^, we hypothesized that hyperactivation of TORC1 may prevent upregulation of *TRA1* upon accumulation of misfolded polyQ expansion. Indeed, *TRA1* was not significantly upregulated after expression of expanded 103Q in the hyperactive *TOR1^L2134M^* strain (Figure 8D). Coupled to the observed decrease in *TRA1* expression upon TORC1 inhibition by rapamycin (Figure 8C), our results establish that TORC1 signaling regulates *TRA1* expression.

**Figure 7:**
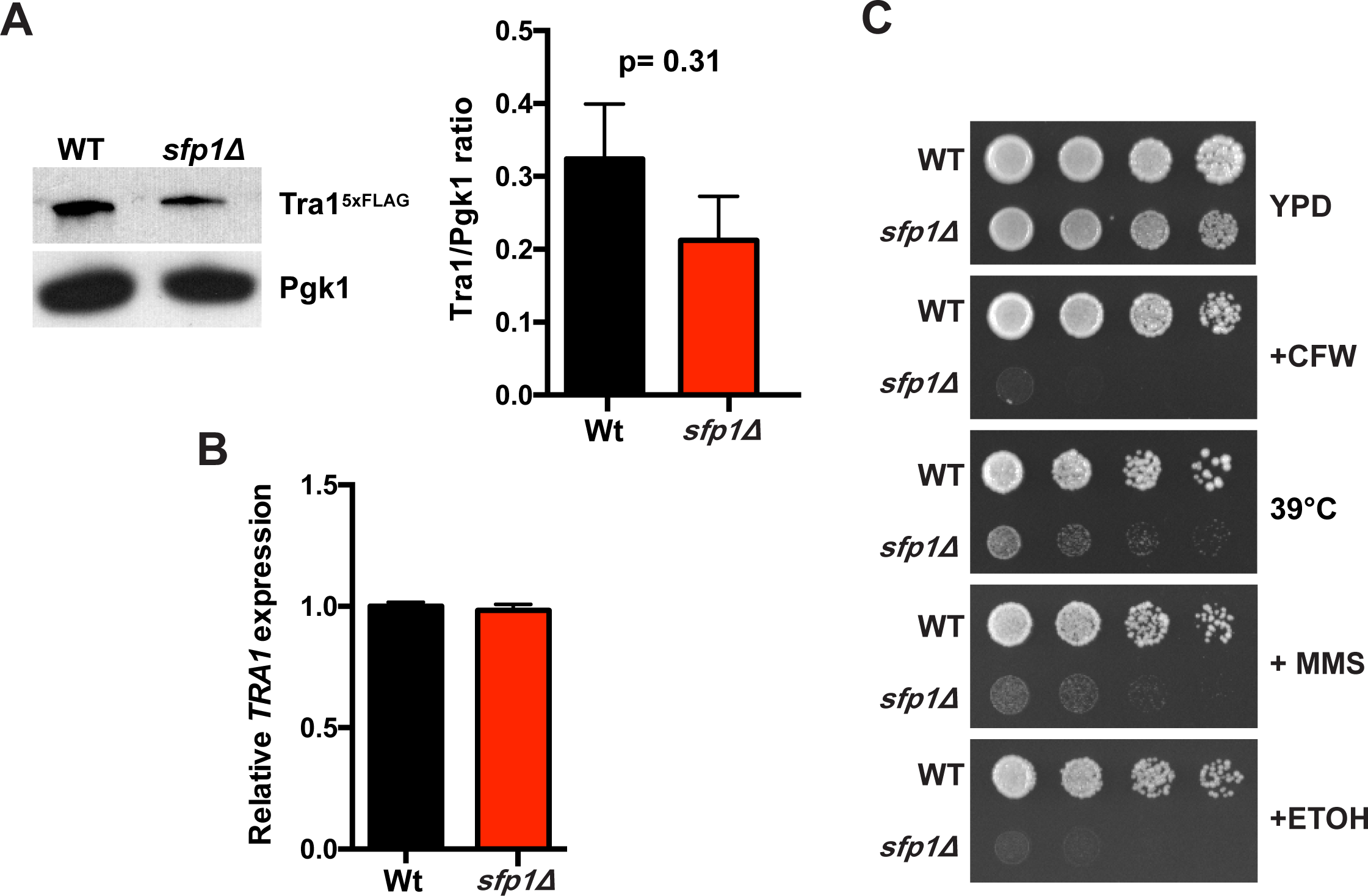
Loss of SFP1 does not significantly affect Tra1 expression but sensitizes cells to stress. **(A)** Deletion of *SFP1* has minimal effect on Tra1 protein abundance. Immunoblot performed with cell lysates of from wild-type and *sfp1Δ* cells expressing a chromosomally integrated Tra1^5x-Flag^. Blot was probed with anti-Flag. Anti-Pgk1 was used as loading control. Densitometric analysis from 3 independent samples is shown. **(B)** Deletion of *SFP1* has minimal effect on *TRA1* mRNA levels. RNA was isolated from wild-type and *sfp1Δ* cells and processed for RT-qPCR to assess transcript levels of *TRA1*. (**C**) Wild-type and *sfp1Δ* cells were spotted on YPD agar plates untreated, supplemented with 6µg/ml calcofluor white (CFW), 5% ethanol (ETOH), 0.03% methyl methanesulfonate (MMS) or incubated at 39°C.

**Figure 8:**
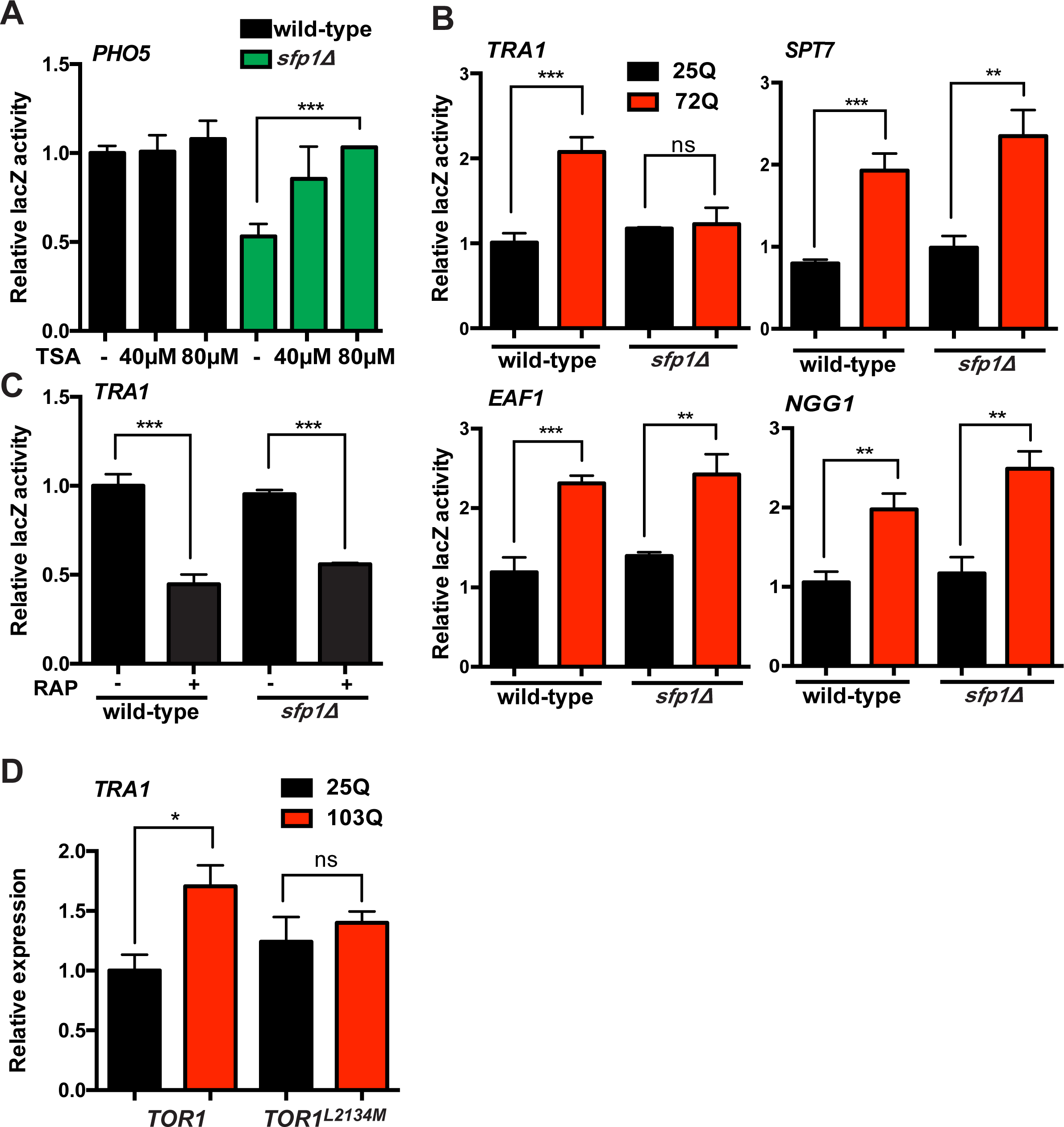
Sfp1 and TORC1 regulate TRA1 expression in response to misfolded polyQ. **(A)** Sfp1 regulates *PHO5* expression. Cells expressing the *PHO5*-LacZ reporter were incubated in absence of phosphate overnight and treated with trichostatin A (TSA) at the indicated concentration. β-galactosidase activity is shown as the average of 3 replicates with the SEM indicated by the error bars. (**B**) Deletion of *SFP1* specifically abolishes *TRA1* upregulation by polyQ. Gene expression from *TRA1* and other SAGA (*NGG1*, *SPT7*) and NuA4 (*EAF1*) promoters after overnight induction of 72Q-CFP compared to 25Q-CFP in wild-type and *sfp1Δ* cells. Gene expression was analyzed using LacZ transcriptional reporters. **p<0.005, ***p<0.001 n=3±SEM. **(C)** Rapamycin treatment decreases transcription from the *TRA1* promoter in both wild-type and *sfp1Δ* cells. Gene expression was analyzed using the *TRA1*-LacZ transcriptional reporter. **p<0.005, ***p<0.001 n=3±SEM. **(D)** Hyperactive TORC1 signaling prevents upregulation of *TRA1* by polyQ. RNA was isolated from wild-type and *TOR1^L2134M^* cells after overnight induction of 25 and 103Q Htt^ex1^-ymsfGFP in galactose media and processed for RT-qPCR to assess transcript levels of *TRA1*.

## DISCUSSION

### Tra1/SAGA and polyQ toxicity

The ability of cytoplasmic polyQ expanded Htt^ex1^ to disrupt Tra1 functions echoes previous reports showing that expression of nuclear-targeted Htt^ex1^ results in transcriptional changes similar to deleting components of the SAGA complex^46^. Other reports show a similar phenotype using a different disease-associated polyQ protein, the SAGA-associated spinocerebellar ataxia 7 (SCA7) ^40,46^. We do not detect significant nuclear localization of Htt^ex1^ in our model (Figure 1). Since Tra1 remains localized to the nucleus upon expression of expanded polyQ, it is unlikely that the two directly interact, thus favoring indirect regulation of Tra1 function by polyQ. Our previous genetic screen using a *tra1* mutant allele revealed that Tra1 is linked to cellular responses of proteotoxic stress, such as the heat shock response, mitochondria homeostasis and TORC1 signaling^47^. Interestingly, all these processes are linked to polyQ toxicity^67,69–71,83–90^. It is reasonable to think that impaired Tra1-regulated transcription has an important role in the toxic phenotype observed in HD models. Indeed, reduced histone acetylation is closely associated with HD and histone deacetylase inhibitors improve the HD phenotype in animal models^23,24,26,27,91,92^. Thus, better characterization of the role of the major regulators of chromatin remodeling and gene expression and their targets provides insight into how modulating acetylation in HD can be beneficial.

### Sfp1 regulates *TRA1* expression in presence of toxic polyQ proteins

Sfp1 is well characterized for its role in adjusting cell size and ribosome production during stress^53–55,76,77^, yet its functions outside the control of ribosome biogenesis are understudied. Recently, Matthew *et al.* found that sequestration of the splicing factor Hsh155 during genotoxic stress was regulated via TORC1 signaling through Sfp1^93^. Together with our data showing that Sfp1 regulates *TRA1* expression, this study supports an expanded role for Sfp1 in stress response, beyond regulating ribosome biogenesis. The mechanism by which Sfp1 contributes to *TRA1* expression during proteotoxic stress is unclear, but the results mimic our previous findings showing that compromising Tra1 function by mutating its PI3-kinase domain increases transcription of *TRA1*^59^. It appears that the cells use a transcriptional feedback mechanism to increase the expression of Tra1 and other components of the SAGA and NuA4 complexes to compensate for the loss of Tra1 function. Whether a common transcription factor is involved remains unknown.

We show that polyQ expansions affect Tra1 function and increase expression of *TRA1* (Figure 4). PolyQ expansions and expression of the *tra1_Q3_* mutant allele^59^ are the only experimental conditions identified to date that increase *TRA1* expression. Interestingly, polyQ-induced *TRA1* expression requires Sfp1 (Figure 8). Previous microarray analysis revealed that overexpressing *SFP1* does not upregulate *TRA1,* indicating that Sfp1 might regulate Tra1 indirectly^55^. Similarly to what we observed with *TRA1*, deleting *SFP1* abolishes upregulation of the proteasome regulator *ADC17* during tunicamycin-induced endoplasmic reticulum stress^82^. Since in both cases, stress leads to Sfp1 relocalization to the cytoplasm, it appears that Sfp1 regulates these targets indirectly, potentially via its regulation of TORC1. The simplest explanation for *TRA1* transcriptional regulation by Sfp1/TORC1 is that *TRA1* is regulated at multiple levels, one that involves TORC1 but also other pathways modulated by toxic polyQ expansions. Finally, whereas there is no homologue of *SFP1* in mammals, Sfp1 shares key functions with the mammalian proto-oncogene *MYC*. These functions include regulating ribosomal protein gene expression and cell growth^94^. Importantly, mammalian TRRAP regulates Myc functions^95–98^. These results delineate a link between Myc and TRRAP that is reminiscent of the Sfp1/Tra1 connection observed in yeast.

### TORC1, SAGA and polyQ toxicity

Our data show that TORC1 has to be finely tuned for yeast cells to compensate for polyQ toxicity. This is not surprising considering the multitude of cellular processes regulated by TORC1^99–101^. Interestingly, both activation and inhibition of TORC1 have been reported to be protective in rodent models of HD^67,69–71^. Our results suggest that downstream targets of TORC1 function in mammalian cells, similar to Sfp1 in yeast, to regulate polyQ toxicity. The inter-relationship of these targets is important for understanding the therapeutic potential of TORC1 for HD. A *TRA1* mutant allele is hypersensitive to rapamycin and results in a synthetic slow growth phenotype with *tor1* deletion^47^. In fission yeast, SAGA control of cell proliferation and differentiation in response to starvation requires the differential TORC1/TORC2-dependent phosphorylation of the SAGA component Taf12^102^. TORC1 also regulates yeast histone acetylation by regulating the sirtuin deacetylases Hst3 and Hst4^103^ and integrates signals through the INO80 chromatin remodeling complex^104^. Because of the breath of cellular functions affected by TORC1 and transcriptional regulators like SAGA, a comprehensive understanding on how these pathways converge to regulate homeostasis both in normal and disease states is crucial. Inhibiting histone deacetylase is protective in various disease models, including HD^91,92,105,106^. Defining how disease-associated proteins affect the global acetylome and identify specific targets for acetylation, including non-histone targets, could lay the basis for the development of new small molecule/compounds for combinational therapy.

## ACKNOWLEDGEMENTS

This study is supported by an operating grant from the Canadian Institute for Health Research (CIHR) to MLD and PL. CJB is supported by a Discovery grant (RGPIN-2015-04394) from the Natural Sciences and Engineering Research Council of Canada (NSERC). YJ is supported by an MSc to PhD Transfer Scholarship from the Schulich Faculty of Medicine and Dentistry at The University of Western Ontario. MDB was supported by an NSERC Alexander Graham Bell Canada Graduate Scholarship and an Ontario Graduate Scholarship and now holds an NSERC Doctoral Scholarship. KA is supported by a Canada Graduate Scholarship from NSERC. The authors thank Dr Tatsuya Maeda (University of Tokyo) for the *TOR1* mutant strain.

---

**Supplemental Figure 1:**
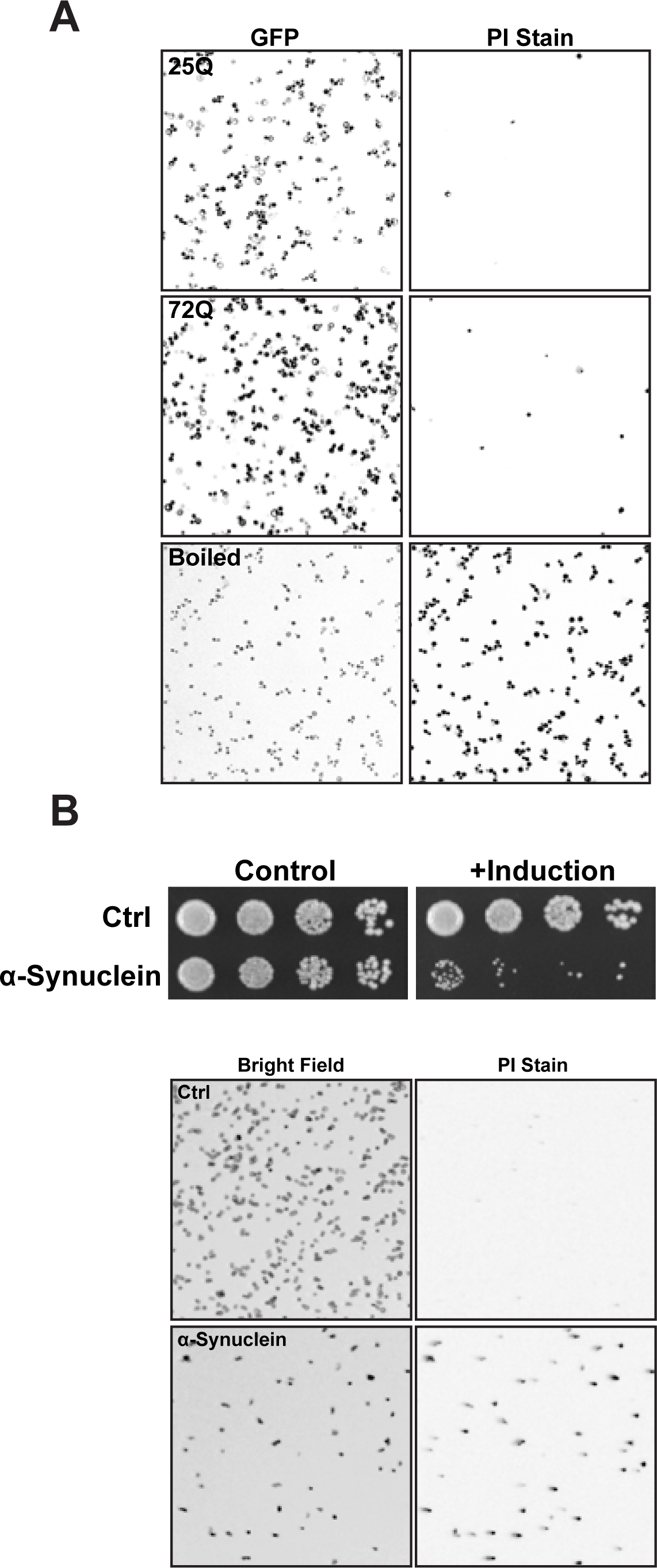
PolyQ expansions do not cause cell death. **(A)** Representative fluorescent images of wild-type cells expressing 25Q or 72Q Htt^ex1^-CFP after overnight induction in galactose containing media and stained with propidium iodide (PI). Boiled cells are shown as a positive control. **(B)** α-synuclein expression results in yeast cell death. Growth of cells expressing an empty vector or - synuclein-GFP was assessed by serial dilutions on SC plates containing either glucose (control) or galactose (induced). Staining with PI assessed cell viability.

**Supplemental Figure 2:**
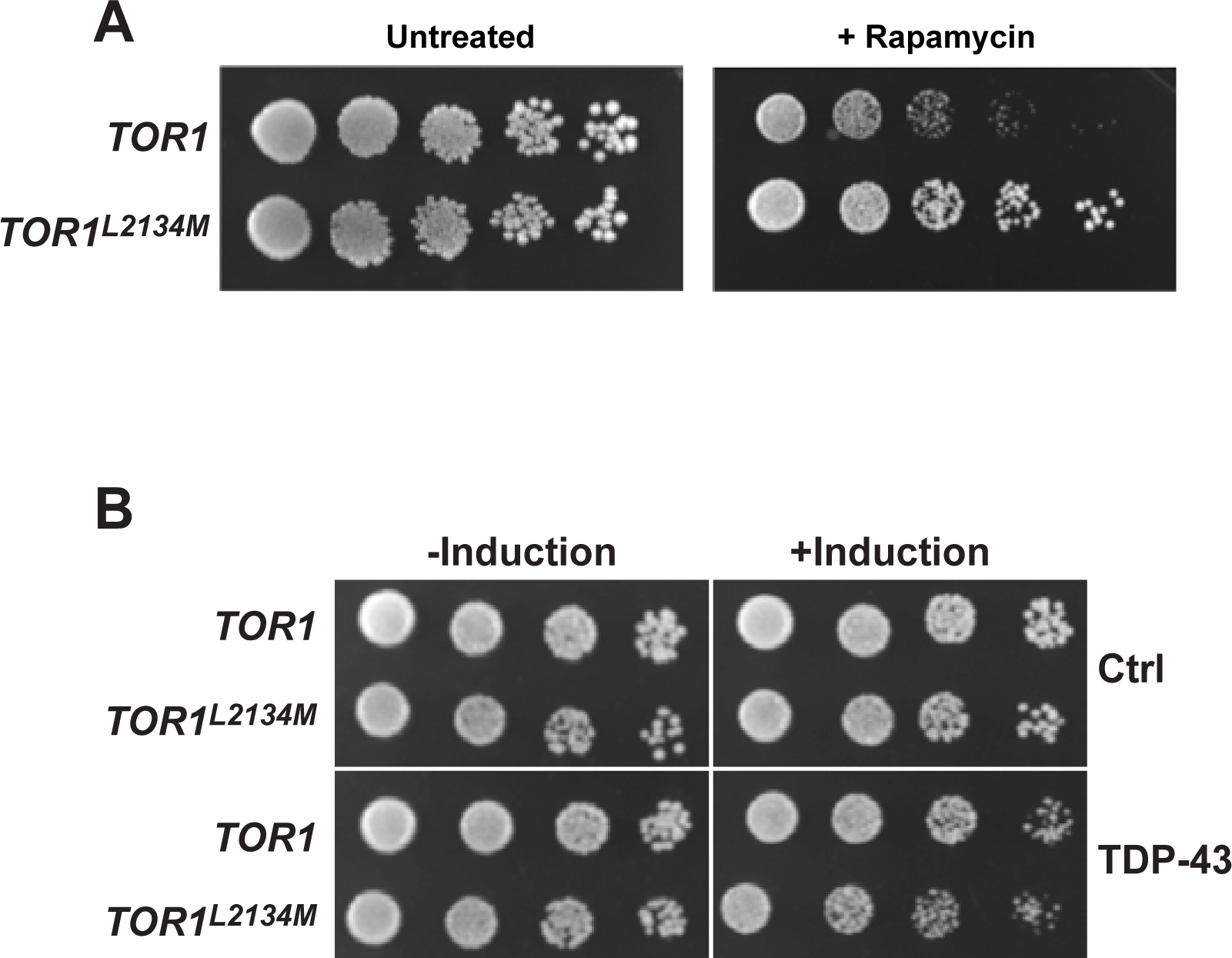
Hyperactive TORC1 signaling has minimal effect on TDP-43 toxicity. **(A)** Growth of wild-type and *TOR1^L2134M^* cells on agar plate containing 10 ng/ml rapamycin **(B)** Growth of wild-type and *TOR1^L2134M^* cells expressing an empty vector or TDP-43 was assessed by serial dilutions on SC plates containing either glucose (control) or galactose (induced).

## References

1. Klaips, C. L., Jayaraj, G. G. & Hartl, F. U. Pathways of cellular proteostasis in aging and disease. J Cell Biol 217, 51–63 (2018).

2. Balch, W. E., Morimoto, R. I., Dillin, A. & Kelly, J. W. Adapting proteostasis for disease intervention. Science 319, 916–919 (2008).

3. Henning, R. H. & Brundel, B. J. J. M. Proteostasis in cardiac health and disease. Nat Rev Cardiol 14, 637–653 (2017).

4. Ruegsegger, C. & Saxena, S. Proteostasis impairment in ALS. Brain Res 1648, 571–579 (2016).

5. Huang, C. C. et al. Amyloid formation by mutant huntingtin: threshold, progressivity and recruitment of normal polyglutamine proteins. Somat Cell Mol Genet 24, 217–233 (1998).

6. Gusella, J. F., MacDonald, M. E. & Lee, J.-M. Genetic modifiers of Huntington’s disease. Mov Disord 29, 1359–1365 (2014).

7. Somatic and gonadal mosaicism of the Huntington disease gene CAG repeat in brain and sperm. J Theilmann - Nature … (1994).

8. Becanovic, K. et al. Transcriptional changes in Huntington disease identified using genome-wide expression profiling and cross-platform analysis. Hum Mol Genet 19, 1438–1452 (2010).

9. Neueder, A. & Bates, G. P. A common gene expression signature in Huntington’s disease patient brain regions. BMC Med Genomics 7, 60 (2014).

10. Tauber, E. et al. Functional gene expression profiling in yeast implicates translational dysfunction in mutant huntingtin toxicity. J Biol Chem 286, 410–419 (2011).

11. Vidal, R. L. et al. Targeting the UPR transcription factor XBP1 protects against Huntington’s disease through the regulation of FoxO1 and autophagy. Hum Mol Genet 21, 2245–2262 (2012).

12. Lajoie, P. & Snapp, E. L. Changes in BiP availability reveal hypersensitivity to acute endoplasmic reticulum stress in cells expressing mutant huntingtin. J Cell Sci 124, 3332–3343 (2011).

13. Vidal, R. L. & Hetz, C. Crosstalk between the UPR and autophagy pathway contributes to handling cellular stress in neurodegenerative disease. Autophagy 8, 970–972 (2012).

14. Duennwald, M. L. & Lindquist, S. Impaired ERAD and ER stress are early and specific events in polyglutamine toxicity. Genes Dev 22, 3308–3319 (2008).

15. Haynes, C. M., Titus, E. A. & Cooper, A. A. Degradation of misfolded proteins prevents ER-derived oxidative stress and cell death. Mol Cell 15, 767–776 (2004).

16. Leitman, J. et al. ER stress-induced eIF2-alpha phosphorylation underlies sensitivity of striatal neurons to pathogenic huntingtin. PLoS ONE 9, e90803 (2014).

17. Walter, P. & Ron, D. The unfolded protein response: from stress pathway to homeostatic regulation. Science 334, 1081–1086 (2011).

18. Verghese, J., Abrams, J., Wang, Y. & Morano, K. A. Biology of the heat shock response and protein chaperones: budding yeast (Saccharomyces cerevisiae) as a model system. Microbiol Mol Biol Rev 76, 115–158 (2012).

19. Chafekar, S. M. & Duennwald, M. L. Impaired heat shock response in cells expressing full-length polyglutamine-expanded huntingtin. PLoS ONE 7, e37929 (2012).

20. Bersuker, K., Hipp, M. S., Calamini, B., Morimoto, R. I. & Kopito, R. R. Heat shock response activation exacerbates inclusion body formation in a cellular model of Huntington disease. J Biol Chem 288, 23633–23638 (2013).

21. Fujikake, N. et al. Heat shock transcription factor 1-activating compounds suppress polyglutamine-induced neurodegeneration through induction of multiple molecular chaperones. J Biol Chem 283, 26188–26197 (2008).

22. Dong, X. et al. The role of h3k4me3 in transcriptional regulation is altered in huntington’s disease. PLoS ONE 10, e0144398 (2015).

23. McFarland, K. N. et al. Genome-wide histone acetylation is altered in a transgenic mouse model of Huntington’s disease. PLoS ONE 7, e41423 (2012).

24. Sharma, S. & Taliyan, R. Transcriptional dysregulation in Huntington’s disease: The role of histone deacetylases. Pharmacol Res 100, 157–169 (2015).

25. Valor, L. M., Guiretti, D., Lopez-Atalaya, J. P. & Barco, A. Genomic landscape of transcriptional and epigenetic dysregulation in early onset polyglutamine disease. J Neurosci 33, 10471–10482 (2013).

26. Guiretti, D. et al. Specific promoter deacetylation of histone H3 is conserved across mouse models of Huntington’s disease in the absence of bulk changes. Neurobiol Dis 89, 190–201 (2016).

27. Steffan, J. S. et al. Histone deacetylase inhibitors arrest polyglutamine-dependent neurodegeneration in Drosophila. Nature 413, 739–743 (2001).

28. Grant, P. A. et al. Yeast Gcn5 functions in two multisubunit complexes to acetylate nucleosomal histones: characterization of an Ada complex and the SAGA (Spt/Ada) complex. Genes Dev 11, 1640–1650 (1997).

29. Allard, S. et al. NuA4, an essential transcription adaptor/histone H4 acetyltransferase complex containing Esa1p and the ATM-related cofactor Tra1p. EMBO J 18, 5108–5119 (1999).

30. Auger, A. et al. Eaf1 is the platform for NuA4 molecular assembly that evolutionarily links chromatin acetylation to ATP-dependent exchange of histone H2A variants. Mol Cell Biol 28, 2257–2270 (2008).

31. Ikura, T. et al. Involvement of the TIP60 histone acetylase complex in DNA repair and apoptosis. Cell 102, 463–473 (2000).

32. McMahon, S. B., Van Buskirk, H. A., Dugan, K. A., Copeland, T. D. & Cole, M. D. The novel ATM-related protein TRRAP is an essential cofactor for the c-Myc and E2F oncoproteins. Cell 94, 363–374 (1998).

33. Baretić, D. & Williams, R. L. The structural basis for mTOR function. Semin Cell Dev Biol 36, 91–101 (2014).

34. Brown, C. E. et al. Recruitment of HAT complexes by direct activator interactions with the ATM-related Tra1 subunit. Science 292, 2333–2337 (2001).

35. Bhaumik, S. R., Raha, T., Aiello, D. P. & Green, M. R. In vivo target of a transcriptional activator revealed by fluorescence resonance energy transfer. Genes Dev 18, 333–343 (2004).

36. Fishburn, J., Mohibullah, N. & Hahn, S. Function of a eukaryotic transcription activator during the transcription cycle. Mol Cell 18, 369–378 (2005).

37. Reeves, W. M. & Hahn, S. Targets of the Gal4 transcription activator in functional transcription complexes. Mol Cell Biol 25, 9092–9102 (2005).

38. Mutiu, A. I. et al. Structure/function analysis of the phosphatidylinositol-3-kinase domain of yeast tra1. Genetics 177, 151–166 (2007).

39. Yang, H. et al. Aggregation of Polyglutamine-expanded Ataxin 7 Protein Specifically Sequesters Ubiquitin-specific Protease 22 and Deteriorates Its Deubiquitinating Function in the Spt-Ada-Gcn5-Acetyltransferase (SAGA) Complex. J Biol Chem 290, 21996–22004 (2015).

40. McMahon, S. J., Pray-Grant, M. G., Schieltz, D., Yates, J. R. & Grant, P. A. Polyglutamine-expanded spinocerebellar ataxia-7 protein disrupts normal SAGA and SLIK histone acetyltransferase activity. Proc Natl Acad Sci U S A 102, 8478–8482 (2005).

41. Burke, T. L., Miller, J. L. & Grant, P. A. Direct inhibition of Gcn5 protein catalytic activity by polyglutamine-expanded ataxin-7. J Biol Chem 288, 34266–34275 (2013).

42. Chen, Y. C. et al. Gcn5 loss-of-function accelerates cerebellar and retinal degeneration in a SCA7 mouse model. Hum Mol Genet 21, 394–405 (2012).

43. Helmlinger, D. et al. Ataxin-7 is a subunit of GCN5 histone acetyltransferase-containing complexes. Hum Mol Genet 13, 1257–1265 (2004).

44. Lan, X. et al. Poly(q) expansions in ATXN7 affect solubility but not activity of the SAGA deubiquitinating module. Mol Cell Biol 35, 1777–1787 (2015).

45. Holmberg, M. et al. Spinocerebellar ataxia type 7 (SCA7): a neurodegenerative disorder with neuronal intranuclear inclusions. Hum Mol Genet 7, 913–918 (1998).

46. Hughes, R. E. et al. Altered transcription in yeast expressing expanded polyglutamine. Proc Natl Acad Sci U S A 98, 13201–13206 (2001).

47. Hoke, S. M. T., Guzzo, J., Andrews, B. & Brandl, C. J. Systematic genetic array analysis links the Saccharomyces cerevisiae SAGA/SLIK and NuA4 component Tra1 to multiple cellular processes. BMC Genet 9, 46 (2008).

48. Meriin, A. B. et al. Huntington toxicity in yeast model depends on polyglutamine aggregation mediated by a prion-like protein Rnq1. J Cell Biol 157, 997–1004 (2002).

49. Duennwald, M. L. Yeast as a platform to explore polyglutamine toxicity and aggregation. Methods Mol Biol 1017, 153–161 (2013).

50. Duennwald, M. L., Jagadish, S., Muchowski, P. J. & Lindquist, S. Flanking sequences profoundly alter polyglutamine toxicity in yeast. Proc Natl Acad Sci U S A 103, 11045–11050 (2006).

51. Duennwald, M. L., Jagadish, S., Giorgini, F., Muchowski, P. J. & Lindquist, S. A network of protein interactions determines polyglutamine toxicity. Proc Natl Acad Sci U S A 103, 11051–11056 (2006).

52. Jiang, Y., Di Gregorio, S. E., Duennwald, M. L. & Lajoie, P. Polyglutamine toxicity in yeast uncovers phenotypic variations between different fluorescent protein fusions. Traffic 18, 58–70 (2017).

53. Fingerman, I., Nagaraj, V., Norris, D. & Vershon, A. K. Sfp1 plays a key role in yeast ribosome biogenesis. Eukaryotic Cell 2, 1061–1068 (2003).

54. Marion, R. M. et al. Sfp1 is a stress- and nutrient-sensitive regulator of ribosomal protein gene expression. Proc Natl Acad Sci U S A 101, 14315–14322 (2004).

55. Jorgensen, P. et al. A dynamic transcriptional network communicates growth potential to ribosome synthesis and critical cell size. Genes Dev 18, 2491–2505 (2004).

56. Janke, C. et al. A versatile toolbox for PCR-based tagging of yeast genes: new fluorescent proteins, more markers and promoter substitution cassettes. Yeast 21, 947–962 (2004).

57. Gietz, R. D. & Woods, R. A. Transformation of yeast by lithium acetate/single-stranded carrier DNA/polyethylene glycol method. Meth Enzymol 350, 87–96 (2002).

58. Mutiu, A. I., Hoke, S. M. T., Genereaux, J., Liang, G. & Brandl, C. J. The role of histone ubiquitylation and deubiquitylation in gene expression as determined by the analysis of an HTB1(K123R) Saccharomyces cerevisiae strain. Mol Genet Genomics 277, 491–506 (2007).

59. Berg, M. D., Genereaux, J., Karagiannis, J. & Brandl, C. J. The Pseudokinase Domain ofSaccharomyces cerevisiaeTra1 Is Required for Nuclear Localization and Incorporation into the SAGA and NuA4 Complexes. G3 (Bethesda) (2018). doi:10.1534/g3.118.200288

60. Brandl, C. J., Furlanetto, A. M., Martens, J. A. & Hamilton, K. S. Characterization of NGG1, a novel yeast gene required for glucose repression of GAL4p-regulated transcription. EMBO J 12, 5255–5265 (1993).

61. Keppler-Ross, S., Noffz, C. & Dean, N. A new purple fluorescent color marker for genetic studies in Saccharomyces cerevisiae and Candida albicans. Genetics 179, 705–710 (2008).

62. Genereaux, J. et al. Genetic evidence links the ASTRA protein chaperone component Tti2 to the SAGA transcription factor Tra1. Genetics 191, 765–780 (2012).

63. Schindelin, J., Rueden, C. T., Hiner, M. C. & Eliceiri, K. W. The ImageJ ecosystem: An open platform for biomedical image analysis. Mol Reprod Dev 82, 518–529 (2015).

64. Kushnirov, V. V. Rapid and reliable protein extraction from yeast. Yeast 16, 857–860 (2000).

65. Hoke, S. M. T. et al. Mutational analysis of the C-terminal FATC domain of Saccharomyces cerevisiae Tra1. Curr Genet 56, 447–465 (2010).

66. Renna, M., Jimenez-Sanchez, M., Sarkar, S. & Rubinsztein, D. C. Chemical inducers of autophagy that enhance the clearance of mutant proteins in neurodegenerative diseases. J Biol Chem 285, 11061–11067 (2010).

67. Ravikumar, B. et al. Inhibition of mTOR induces autophagy and reduces toxicity of polyglutamine expansions in fly and mouse models of Huntington disease. Nat Genet 36, 585–595 (2004).

68. Ravikumar, B., Duden, R. & Rubinsztein, D. C. Aggregate-prone proteins with polyglutamine and polyalanine expansions are degraded by autophagy. Hum Mol Genet 11, 1107–1117 (2002).

69. Lee, J. H. et al. Reinstating aberrant mTORC1 activity in Huntington’s disease mice improves disease phenotypes. Neuron 85, 303–315 (2015).

70. Ferrarelli, L. K. Is mTOR a good guy or bad guy in Huntington’s disease? Sci Signal 8, ec26–ec26 (2015).

71. Pryor, W. M. et al. Huntingtin promotes mTORC1 signaling in the pathogenesis of Huntington’s disease. Sci Signal 7, ra103 (2014).

72. Takahara, T. & Maeda, T. Transient sequestration of TORC1 into stress granules during heat stress. Mol Cell 47, 242–252 (2012).

73. Johnson, B. S., McCaffery, J. M., Lindquist, S. & Gitler, A. D. A yeast TDP-43 proteinopathy model: Exploring the molecular determinants of TDP-43 aggregation and cellular toxicity. Proc Natl Acad Sci U S A 105, 6439–6444 (2008).

74. Loewith, R. & Hall, M. N. Target of rapamycin (TOR) in nutrient signaling and growth control. Genetics 189, 1177–1201 (2011).

75. Urban, J. et al. Sch9 is a major target of TORC1 in Saccharomyces cerevisiae. Mol Cell 26, 663–674 (2007).

76. Singh, J. & Tyers, M. A Rab escort protein integrates the secretion system with TOR signaling and ribosome biogenesis. Genes Dev 23, 1944–1958 (2009).

77. Lempiäinen, H. et al. Sfp1 interaction with TORC1 and Mrs6 reveals feedback regulation on TOR signaling. Mol Cell 33, 704–716 (2009).

78. Lopez, A. D. et al. Proteasomal degradation of Sfp1 contributes to the repression of ribosome biogenesis during starvation and is mediated by the proteasome activator Blm10. Mol Biol Cell 22, 528–540 (2011).

79. Xu, Z. & Norris, D. The SFP1 gene product of Saccharomyces cerevisiae regulates G2/M transitions during the mitotic cell cycle and DNA-damage response. Genetics 150, 1419–1428 (1998).

80. Barbaric, S., Reinke, H. & Hörz, W. Multiple mechanistically distinct functions of SAGA at the PHO5 promoter. Mol Cell Biol 23, 3468–3476 (2003).

81. Gregory, P. D. et al. Absence of Gcn5 HAT activity defines a novel state in the opening of chromatin at the PHO5 promoter in yeast. Mol Cell 1, 495–505 (1998).

82. Rousseau, A. & Bertolotti, A. An evolutionarily conserved pathway controls proteasome homeostasis. Nature 536, 184–189 (2016).

83. Wyttenbach, A. et al. Heat shock protein 27 prevents cellular polyglutamine toxicity and suppresses the increase of reactive oxygen species caused by huntingtin. Hum Mol Genet 11, 1137–1151 (2002).

84. Chafekar, S. M. et al. Pharmacological tuning of heat shock protein 70 modulates polyglutamine toxicity and aggregation. ACS Chem Biol 7, 1556–1564 (2012).

85. Sakahira, H., Breuer, P., Hayer-Hartl, M. K. & Hartl, F. U. Molecular chaperones as modulators of polyglutamine protein aggregation and toxicity. Proc Natl Acad Sci U S A 99 Suppl 4, 16412–16418 (2002).

86. King, M. A. et al. Rapamycin inhibits polyglutamine aggregation independently of autophagy by reducing protein synthesis. Mol Pharmacol 73, 1052–1063 (2008).

87. Oliveira, J. M. A. et al. Mitochondrial dysfunction in Huntington’s disease: the bioenergetics of isolated and in situ mitochondria from transgenic mice. J Neurochem 101, 241–249 (2007).

88. Shirendeb, U. et al. Abnormal mitochondrial dynamics, mitochondrial loss and mutant huntingtin oligomers in Huntington’s disease: implications for selective neuronal damage. Hum Mol Genet 20, 1438–1455 (2011).

89. Costa, V. et al. Mitochondrial fission and cristae disruption increase the response of cell models of Huntington’s disease to apoptotic stimuli. EMBO Mol Med 2, 490–503 (2010).

90. Squitieri, F. et al. Severe ultrastructural mitochondrial changes in lymphoblasts homozygous for Huntington disease mutation. Mech Ageing Dev 127, 217–220 (2006).

91. Jia, H. et al. The effects of pharmacological inhibition of histone deacetylase 3 (HDAC3) in huntington’s disease mice. PLoS ONE 11, e0152498 (2016).

92. Yeh, H. H. et al. Histone deacetylase class II and acetylated core histone immunohistochemistry in human brains with Huntington’s disease. Brain Res 1504, 16–24 (2013).

93. Mathew, V. et al. Selective aggregation of the splicing factor Hsh155 suppresses splicing upon genotoxic stress. J Cell Biol 216, 4027–4040 (2017).

94. Dang, C. V. et al. The c-Myc target gene network. Semin Cancer Biol 16, 253–264 (2006).

95. Zhang, N. et al. MYC interacts with the human STAGA coactivator complex via multivalent contacts with the GCN5 and TRRAP subunits. Biochim Biophys Acta 1839, 395–405 (2014).

96. Bhattacharya, S. & Ghosh, M. K. HAUSP regulates c-MYC expression via de-ubiquitination of TRRAP. Cell Oncol (Dordr) 38, 265–277 (2015).

97. Kenneth, N. S. et al. TRRAP and GCN5 are used by c-Myc to activate RNA polymerase III transcription. Proc Natl Acad Sci U S A 104, 14917–14922 (2007).

98. Liu, X., Tesfai, J., Evrard, Y. A., Dent, S. Y. R. & Martinez, E. c-Myc transformation domain recruits the human STAGA complex and requires TRRAP and GCN5 acetylase activity for transcription activation. J Biol Chem 278, 20405–20412 (2003).

99. Betz, C. & Hall, M. N. Where is mTOR and what is it doing there? J Cell Biol 203, 563–574 (2013).

100. Liko, D. & Hall, M. N. mTOR in health and in sickness. J Mol Med 93, 1061–1073 (2015).

101. González, A. & Hall, M. N. Nutrient sensing and TOR signaling in yeast and mammals. EMBO J 36, 397–408 (2017).

102. Laboucarié, T. et al. TORC1 and TORC2 converge to regulate the SAGA co-activator in response to nutrient availability. EMBO Rep 18, 2197–2218 (2017).

103. Workman, J. J., Chen, H. & Laribee, R. N. Saccharomyces cerevisiae TORC1 Controls Histone Acetylation by Signaling Through the Sit4/PP6 Phosphatase to Regulate Sirtuin Deacetylase Nuclear Accumulation. Genetics 203, 1733–1746 (2016).

104. Beckwith, S. L. et al. The INO80 chromatin remodeler sustains metabolic stability by promoting TOR signaling and regulating histone acetylation. PLoS Genet 14, e1007216 (2018).

105. Valor, L. M. & Guiretti, D. What’s wrong with epigenetics in Huntington's disease? Neuropharmacology 80, 103–114 (2014).

106. Valor, L. M. Epigenetic-based therapies in the preclinical and clinical treatment of Huntington’s disease. Int J Biochem Cell Biol 67, 45–48 (2015).

